# Auditory prediction errors and auditory white matter microstructure as predictors of psychotic experiences in healthy individuals

**DOI:** 10.1101/544452

**Authors:** R. Randeniya, L.K.L. Oestreich, M.I. Garrido

## Abstract

Our sensory systems actively predict sensory information based on previously learnt patterns. An inability to accurately predict forthcoming information results in prediction errors. Individuals with schizophrenia consistently show reduced auditory prediction errors as well as reduced microstructure in auditory white matter pathways. However, it is not clear if also healthy individuals with psychotic experiences demonstrate such deficits. Participants underwent electroencephalography (EEG) recordings while listening to a simple two-tone duration deviant oddball paradigm (N=103) and a stochastic oddball paradigm (N=89). A subset of participants (N=89) also underwent diffusion-weighted magnetic resonance imaging (MRI), from which fractional anisotropy (FA), a measure of overall white matter microstructure, was obtained for auditory pathways namely the auditory interhemispheric pathway, as well as the left and right arcuate fasciculi. We investigated both structural and functional predictors of positive psychotic experiences in healthy participants as measured by the Community Assessment for Psychic Experiences positive dimension (CAPE+) scores. Prediction errors evoked by the classical oddball paradigm failed to reveal significant effects, whereas the stochastic oddball paradigm revealed significant clusters at typical mismatch negativity periods predictive of CAPE+ scores. Furthermore, we show that white matter microstructure from auditory pathways in addition to mismatches significantly predict CAPE+ scores. We suggest that structural and functional prediction error measures together may have potential in predicting psychotic experiences in the healthy population.

## 1. Introduction

Auditory prediction errors, particularly measured with electroencephalography (EEG), have gained headway in the search for biomarkers of schizophrenia. Mismatch negativity (MMN) is a component of prediction errors, evoked by stimuli that differ from a learnt pattern (Belger et al., 2012; Nagai et al., 2013b; Näätänen, 2014). MMN has been suggested as a potential biomarker for schizophrenia (Nagai et al., 2013a) due to the fact that patients with schizophrenia consistently exhibit a robust attenuation of the auditory MMN (Todd et al., 2008; Horton et al., 2011; Erickson et al., 2016). Furthermore, MMN has also been shown to be attenuated in first-episode psychosis (Haigh et al., 2016a; Salisbury et al., 2016) and individuals at high-risk for schizophrenia (Atkinson et al., 2012; Shaikh et al., 2012; Nagai et al., 2013b; Perez et al., 2014; Solis-Vivanco et al., 2014). Critically, studies have shown that the MMN response predicts the development of schizophrenia in individuals at high-risk (Bodatsch et al., 2011; Bodatsch et al., 2015). The P300, a positive peak occurring 250-400ms after tone onset (Sur and Sinha, 2009), is a later component of the prediction error response elicited with auditory oddball paradigms. The P300 component has been shown to be reduced in chronic schizophrenia (Ford et al., 2001; Winterer et al., 2003) and in first-episode schizophrenia (Qiu et al., 2014).

Findings in patients with schizophrenia can often be confounded by disease stage and medication use. Schizotypy is a construct that refers to healthy individuals with psychotic-like experiences, such as delusions and/or hallucinations, and is thought to be the sub-clinical manifestation of schizophrenia in the general population (Ettinger et al., 2015). Studying psychotic experiences in healthy individuals has the advantage of avoiding the confounding factors present in many schizophrenia studies (medication, co-morbidities, etc) and could therefore provide insights into the very early precursors of schizophrenia. It has been shown that prediction errors reduce with increasing delusional-like experiences in healthy populations (Corlett, 2012). Furthermore, the trait phenotype social disorganization, which is shared by autism and schizotypy, has been linked to reduced fronto-temporal response to deviant tones in a magnetoencephalography study (Ford et al., 2017).

MMN to duration deviants is a particularly strong candidate biomarker of schizophrenia (Erickson et al., 2016). Other more complex paradigms with double deviants (Perez et al., 2014), tone omission deviants (Salisbury and McCathern, 2016) and pattern violations (Haigh et al., 2016b) have also demonstrated significant attenuation in MMN in patients with schizophrenia and individuals at high-risk. A more recent meta-analysis reported that the magnitude of attenuation in MMN in schizophrenia patients to complex paradigms are as large as that in simple paradigms and therefore suggest that pitch and duration paradigms provide simpler avenues for further research (Avissar, 2018). However, there is limited research investigating a putative association between psychotic experiences in the healthy population and prediction errors in both simple and complex paradigms. It is plausible that more complex oddball paradigms are more sensitive to less severe expressions of psychotic experiences in the general population, a conjecture we set out to test in this study.

Effective connectivity studies of frequency MMN in healthy individuals have identified primary auditory cortex (A1), superior temporal gyrus (STG), and the inferior temporal gyrus (IFG) as key nodes in the generation of prediction errors (Garrido et al., 2009). Frequency MMN paradigms in patients with schizophrenia have shown that patients have reduced connectivity within this network during auditory mismatch responses (Dima et al., 2012; Larsen, 2018). Interestingly, these effectively connected areas are also structurally connected via auditory white matter pathways. One such pathways is the auditory interhemispheric pathway, which is the part of the corpus callosum that connects bilateral A1, and the other is the arcuate fasciculus, an association tract which connects STG and IFG. In schizophrenia, reduced white matter microstructure in the auditory interhemispheric pathway(Wigand et al., 2015) and the arcuate fasciculus(Geoffroy et al., 2014; McCarthy-Jones et al., 2015) have both been linked to auditory verbal hallucinations, which is the most prominent psychotic symptom. In healthy individuals, white matter connectivity reductions in the arcuate fasciculus and the corpus callosum more broadly have been linked to psychotic-like experiences (Nelson et al., 2011; Oestreich et al., 2018).

Based on the robust findings of MMN attenuation in schizophrenia and high-risk individuals who later transition, as well as known white matter reductions in auditory networks subserving prediction error generation, we set out to investigate whether these deficits could also be observed in healthy individuals with varying degrees of psychotic experiences. In order to account for the possibility that a more complex oddball paradigm would be more sensitive to detecting prediction error deficits in healthy individuals with psychotic experiences, we included a stochastic oddball paradigm, whereby deviant tones are outliers within a Gaussian distribution. This was done in addition to a traditional duration deviant paradigm, which has consistently been reported to identify prediction error deficits in schizophrenia. Due to the preliminary supporting findings of MMN attenuation and reduced auditory microstructure associated with schizotypal traits, we hypothesized that the prediction error amplitude would be attenuated as the number of psychotic-like experiences increases and that the microstructure of the arcuate fasciculus and the auditory interhemispheric pathway would predict psychotic experiences in heathy individuals.

## 2. Methods

**Table 1:**
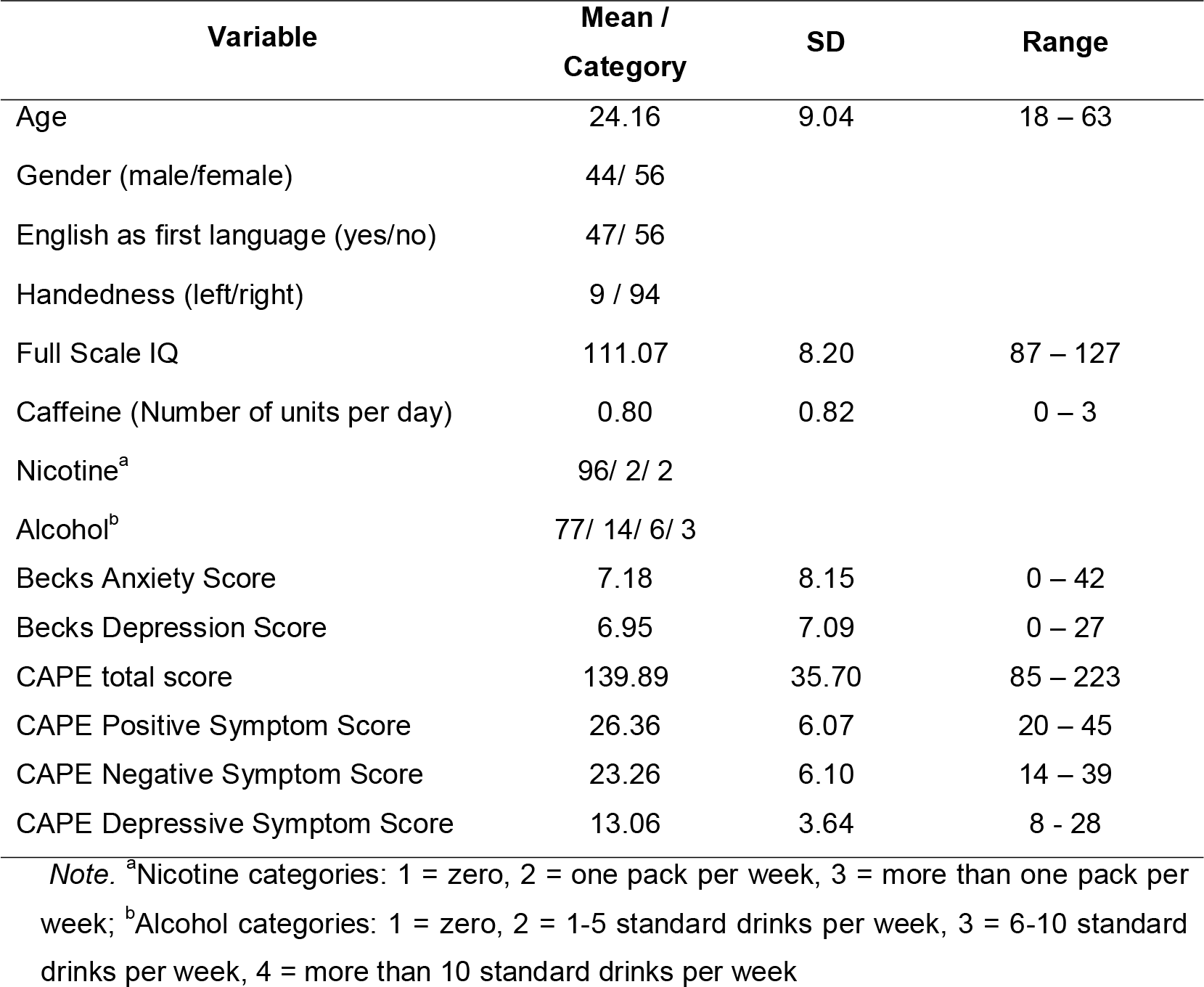
Demographic information

| Variable | Mean /<br>Category | SD | Range |
| --- | --- | --- | --- |
| Age | 24.16 | 9.04 | 18 – 63 |
| Gender (male/female) | 44/ 56 |  |  |
| English as first language (yes/no) | 47/ 56 |  |  |
| Handedness (left/right) | 9 / 94 |  |  |
| Full Scale IQ | 111.07 | 8.20 | 87 – 127 |
| Caffeine (Number of units per day) | 0.80 | 0.82 | 0 – 3 |
| Nicotine<sup>a</sup> | 96/ 2/ 2 |  |  |
| Alcohol<sup>b</sup> | 77/ 14/ 6/ 3 |  |  |
| Becks Anxiety Score | 7.18 | 8.15 | 0 – 42 |
| Becks Depression Score | 6.95 | 7.09 | 0 – 27 |
| CAPE total score | 139.89 | 35.70 | 85 – 223 |
| CAPE Positive Symptom Score | 26.36 | 6.07 | 20 – 45 |
| CAPE Negative Symptom Score | 23.26 | 6.10 | 14 – 39 |
| CAPE Depressive Symptom Score | 13.06 | 3.64 | 8 - 28 |

*Note.* <sup>a</sup>Nicotine categories: 1 = zero, 2 = one pack per week, 3 = more than one pack per week; <sup>b</sup>Alcohol categories: 1 = zero, 2 = 1-5 standard drinks per week, 3 = 6-10 standard drinks per week, 4 = more than 10 standard drinks per week

### 2.1 Participants

One-hundred and three participants were recruited via the University of Queensland (UQ) online recruitment system (SONA) and the UQ newsletter. Participants were between ages 18 and 65 (*M* = 24.67, *SD* = 9.77) and 55.3% (*n* = 57) were female. Participants completed self-report questionnaires including demographic data, Beck’s Anxiety Inventory (Beck et al., 1988), Beck’s Depression Inventory(Beck et al., 1961), and the Community Assessment of Psychic Experience(CAPE)(Stefanis et al., 2002); See distribution of CAPE scores in Figure 1. Descriptive statistical analysis of demographic data and psychometric scales were performed using SPSS(IBM Corp, 2017); (See Table 1). Participants were financially reimbursed for their time. Exclusion criteria included any diagnosis of psychiatric or neurological condition, or head injury with loss of consciousness. Three participants who were taking antidepressant medications during the time of the study were excluded from the analyses to rule out possible medication effects. Ethical clearance for the study was obtained from the Ethics Committee at UQ.

**Figure 1:**
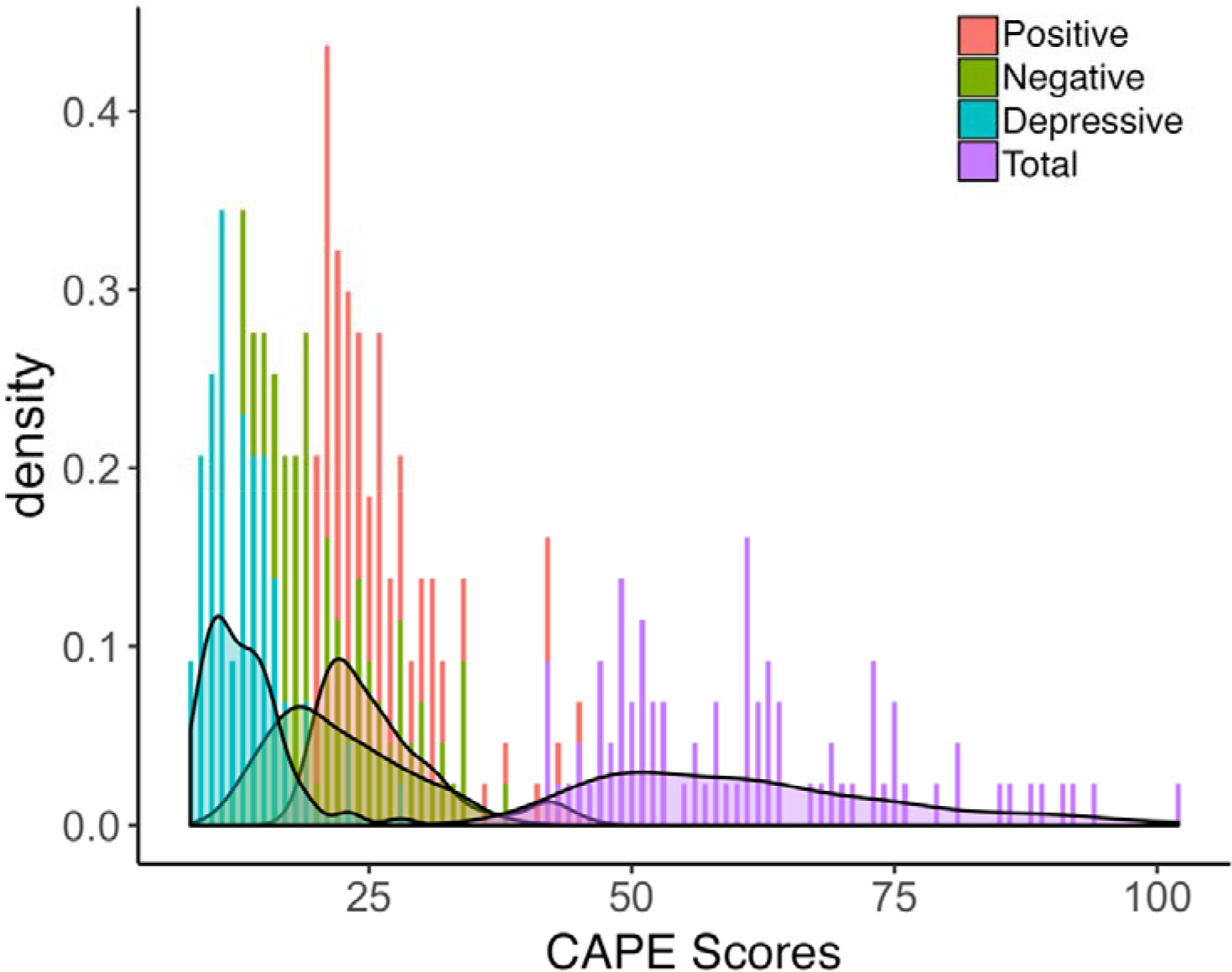
Distribution of CAPE Scores. Participant’s scores on the community assessment of psychic experiences (CAPE-42) questionnaire. Positive(red), negative(green) and depressive(blue) subscale and total (purple) scores shown.

### 2.2 Experimental Procedure

#### 2.2.1 Classical oddball paradigm

All participants underwent a classic auditory duration oddball paradigm which consisted of two blocks. The *Long deviant block* consisted of standard tones (500Hz, 50ms duration, 80% probability), and deviant tones (500Hz, 100ms duration, 20% probability). The *Short deviant block* consisted of the reverse, i.e. standard tones - 500Hz, 100ms duration, 80% probability. All tones occurred at an inter-stimulus interval of 500ms. The order of the blocks was counterbalanced across participants. Participants also engaged in a simultaneous visual letter 1-back task (Miller et al., 2009). All stimuli were written and delivered in MATLAB using Psychtoolbox3(Brainard, 1997; Pelli, 1997). This experimental component lasted for approximately 20 minutes.

#### 2.2.2 Stochastic oddball paradigm

Eighty-nine participants also underwent a stochastic frequency oddball paradigm (Garrido et al., 2013) and a simultaneous 2-back task (Sweet, 2011). Participants listened to a stream of tones with log-frequencies sampled from two Gaussian distributions with equal means (500Hz) and different standard deviations (narrow: *σ*_*n*_ = .5 octaves; broad: *σ*_*b*_ = 1.5 octaves). All tones were played with a duration of 50ms with 10ms smooth rise and fall periods and inter-stimulus intervals of 500ms. 10% of the tones were defined as standard tones, which were played at 500Hz, i.e. the mean of both distributions and 10% of the tones were defined at deviant tones, which were played at 2000Hz, i.e. as outliers to the two distributions. Standard and deviant tones were inserted into the sound stream at random time points. Participants were instructed to disregard the tones and to focus on the visual task instead. This experimental component lasted for approximately 30 minutes and was divided into 4 blocks. The narrow and broad distribution conditions were presented in separate blocks and the order of the blocks was counter-balanced across participants.

### 2.3 EEG data

#### 2.3.1 Acquisition and processing

Throughout the auditory oddball experiments, an electroencephalogram (EEG) was recorded using a 64 electrode-cap with BioSemi ActiView system at a sampling rate of 1024Hz and the following specifications: 417Hz bandwidth (3dB) and 18dB/octave roll-off. Further electrodes were placed on the outer canthi of both eyes, as well as below and above the left eye to measure eye movement. Triggers were marked in the EEG data at the onset of each tone. The EEG raw data was pre-processed using SPM12(Welcome Trust Centre for Neuroimaging, London: http://www.fil.ion.ucl.ac.uk/spm/) run on MATLAB version R2015b (MathWorks). EEG data were segmented into 500ms intervals, consisting of 100ms pre-, and 400ms post-stimulus onset. Topography based correction was used to correct for eye blinks. Data were filtered using a high-pass filter of 0.5Hz and a low-pass filter of 40Hz filter, respectively. Trials containing artefacts with voltages exceeding ±50μV, were rejected. The remaining artefact-free trials in each condition were robustly averaged(Wager et al., 2005) to event-related potentials (ERPs) for each participant and baseline corrected using the pre-stimulus interval.

#### 2.3.2 Spatiotemporal analysis

Using SPM12 ERPs were converted into 3D spatiotemporal volumes for each condition and participant, by interpolating and dividing the scalp data per time point into a 2-dimensional (2D) 32×32 matrix. One 2D image was obtained for each time bin and stacked according to their pre-stimulus temporal order. This resulted in a 3D spatiotemporal image volume with 32×32×81 dimensions per participant. In order to test the effect of prediction, the 3D image volumes were modelled with a full-factorial general linear model using 2×2 ANCOVA designs, with *Deviant Type* (long/short) and *Surprise* (standards/deviants) in the classical oddball paradigm, and *Variance* (narrow/broad) and *Surprise* (standards/deviants) in the stochastic oddball paradigm, as within-subject factors. As it has previously been reported that MMN decreases with age(Cheng et al., 2013), age was included as a covariate. Spatiotemporal maps are visualized using Porthole and Stormcloud toolbox for MATLAB (Taylor and Garrido, 2019).

We then investigated whether healthy individuals with increasing psychotic experiences show reductions in mismatch responses (classical oddball paradigm) and are less able to learn about statistical regularities in a stochastic environment (stochastic oddball paradigm). To this end, we ran regression analyses with individually computed contrast images for the effect of *Surprise* in the Long deviant block of the classical oddball paradigm and for the interaction *Variance*Surprise* in the stochastic oddball paradigm as the outcome variables and CAPE positive dimension scores (CAPE+) as predictor variable were performed. Age was again added as a covariate. In order to isolate the effects of psychotic experiences on the MMN from other measures of psychopathology we added Beck anxiety scores, CAPE depression, CAPE distress and CAPE negative symptom scores as covariates to the analysis. All statistical analyses were corrected for multiple-comparisons using a family-wise error (FWE) rate at an alpha level of 0.05. All *p*-values reported are cluster-FWE corrected.

### 2.4 DWI data

#### 2.4.1 Acquisition and pre-processing

A subsample of 89 (age *M*=24.69; *SD*=10.13) participants also underwent diffusion-weighted and T1-weighted MRI on a 3T Siemens Magnetom TrioTim system. Imaging parameters included TR = 8600ms, TE = 116ms, FOV = 220mm, 2.0 × 2.0 × 2.0mm slice thickness and 15min acquisition time. Diffusion data was acquired at *b*-value=1000s/mm^2^ (32 directions) and *b*-value=3000s/mm^2^ (64 directions). Three interspersed *b*=0 images were obtained including one phase-encoded *b*=0. A T1-weighted image data set was acquired with the MP2RAGE sequence(Marques et al., 2010) with FoV 240mm, 176 slices, 0.9mm isotropic resolution, TR=4000ms, TE=2.92ms, TI1=700ms, TI2=2220ms, first flip angle=6°, second flip angle=7°, and 5min acquisition time.

The DWI volumes were pre-processed using FSL (Functional MRI of the Brain Software Library). Signal intensity inhomogeneities were removed(Zhang et al., 2001). Intra-scan misalignments due to head movements and eddy currents were removed using FSL TOPUP(Smith et al., 2004) and EDDY(Andersson and Sotiropoulos, 2016). Diffusion-weighted images and T1-weighted images were co-registered using boundary-based registration(Greve and Fischl, 2009). Five tissue-type segmentation of T1-weighted images was conducted using the *recon-all* command in Freesurfer (http://surfer.nmr.mgh.harvard.edu/)(Dale et al., 1999). Response functions were estimated using MRtrix (https://github.com/MRtrix3/mrtrix3) multi-shell, multi-tissue algorithm(Jeurissen et al., 2014).

#### 2.4.2 Tractography and Fractional Anisotropy

All tractography steps were undertaken in MRtrix3. Multi-tissue constrained spherical deconvolution was applied to obtain fibre orientation distributions for each voxel (Jeurissen et al., 2014). Probabilistic streamlines of the right arcuate fasciculus, left arcuate fasciculus and auditory interhemispheric tract (Figure 2) were reconstructed using anatomically-constrained tractography(Smith et al., 2015). For each tract, regions of interest (ROI) were manually drawn. The left/ right arcuate fasciculus were seeded from the left/right Broca’s area, with inclusion masks in the respective Geschwind’s territory and target ROIs at Wernicke’s area. The auditory interhemispheric pathway was seeded from the left primary auditory cortex with inclusion masks in the left and right Wernicke’s region and a target ROI in the right primary auditory cortex. Tractography procedure was guided by 500,000 seeds, 4mm minimum and 200mm maximum track length, 1mm step size and maximum number of attempts per seed 500,000. Spherical Deconvolution Informed Filtering of Tractograms (SIFT2) was used to ensure that the reconstructed white matter tracts reflected biologically meaningful connectivity, reducing inadequacies resulting from the reconstruction method(Smith et al., 2015). The fractional anisotropy (FA) for each voxel was then calculated, and the mean FA for each tract was obtained.

**Figure 2:**
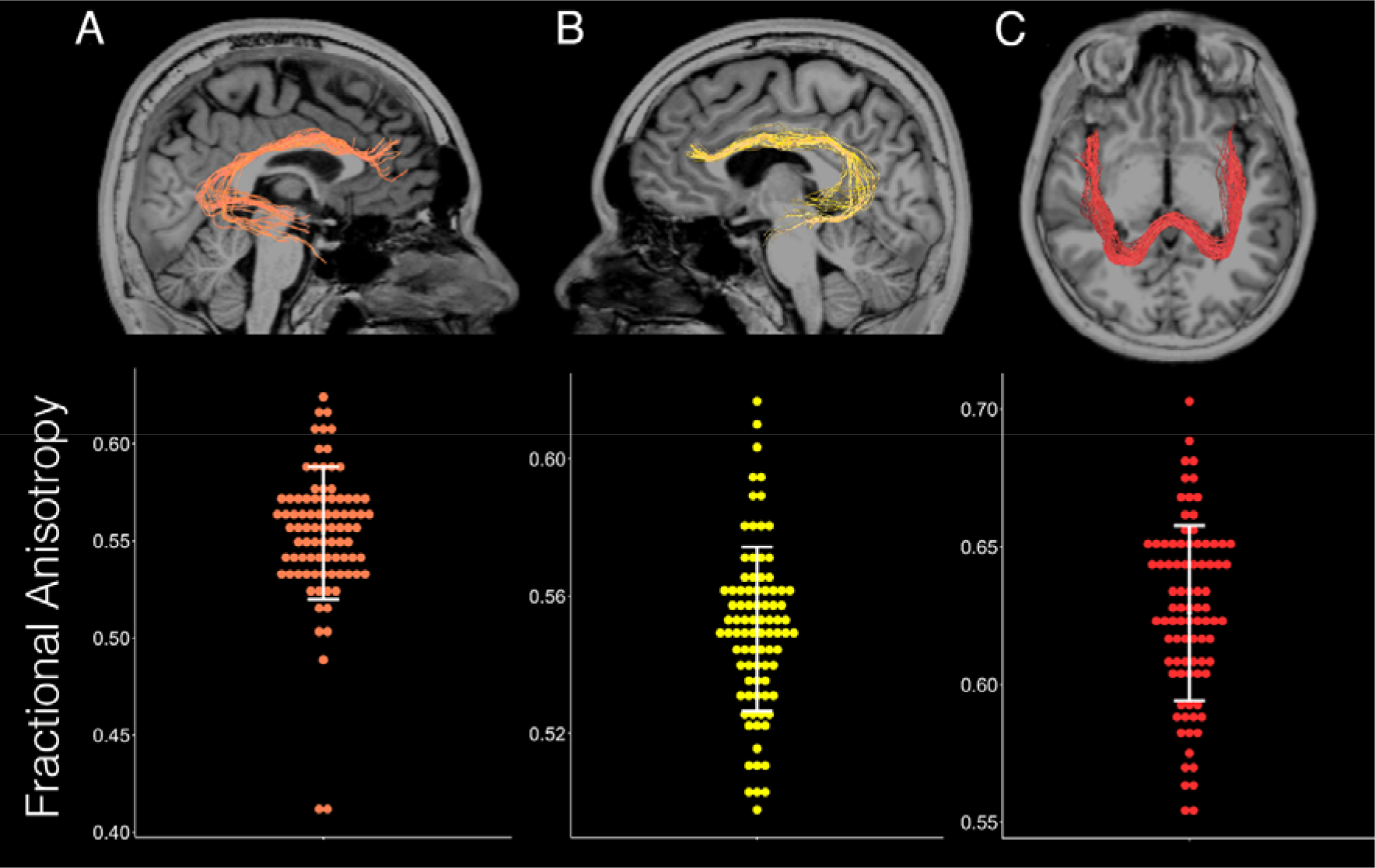
Auditory white matter pathways. (A) The auditory interhemispheric pathway, (B) the right arcuate fasciculus, and (C) the left arcuate fasciculus from three participants. (D) FA of the three white matter pathways for the whole sample (N=89 individuals). Each dot represents one participant and the error bar represents the standard error of the mean.

### 2.5 Regression model for predicting psychotic experiences

A multiple linear regression analysis was conducted to test for functional and structural brain network predictors of psychotic experiences. Field Intensity values for significant clusters from the *surprise*variance* interaction arising from the spatiotemporal analysis were added as predictor variables in a first step. In order to investigate whether estimates of structural connectivity could significantly improve predictions of psychotic experiences, FA of each tract (left arcuate fasciculus, right arcuate fasciculus, auditory interhemispheric pathway) were added as predictors in a second step. CAPE+ scores was the outcome variable.

## 3. Results

In what follows we undertook a spatiotemporal analysis of the entire prediction error response for the whole epoch (400ms), including both MMN and P300.

### 3.1 Spatiotemporal Analysis

#### 3.1.1 Classical Oddball Paradigm

Prediction error waveforms (Deviants > Standards) from a fronto-central channel for each deviant block are shown in Figure 3A and 3B. We conducted a spatiotemporal analysis to investigate if there was an effect of surprise and deviant type. An ANCOVA revealed significant main effects of *Surprise* and *Deviant Type* and a significant interaction of *Surprise*Devian*t type (Figure 3). The main effect of *Surprise* (Figure 3C) showed significant clusters over frontocentral channels at 235ms (*F* = 474.17, *p* <0.001), over temporoparietal channels at 395ms (*F* = 72.23, *p* <0.001) and 400ms (*F* = 73.40, *p* <0.001). Greater prediction errors for long than short duration deviants are demonstrated by the *Surprise*Deviant Type* (Figure 3D) interaction, which showed significant clusters beginning at 180ms (*F* = 327.78, *p* <0.001) over frontocentral channels, at 280ms temporoparietal channels (*F* = 27.91), *p* <0.001) and at 400ms (*F* = 27.69, *p* <0.001) over temporo-parietal channels. This finding is in line with studies with oddball paradigms(Peter et al., 2010).

**Figure 3:**
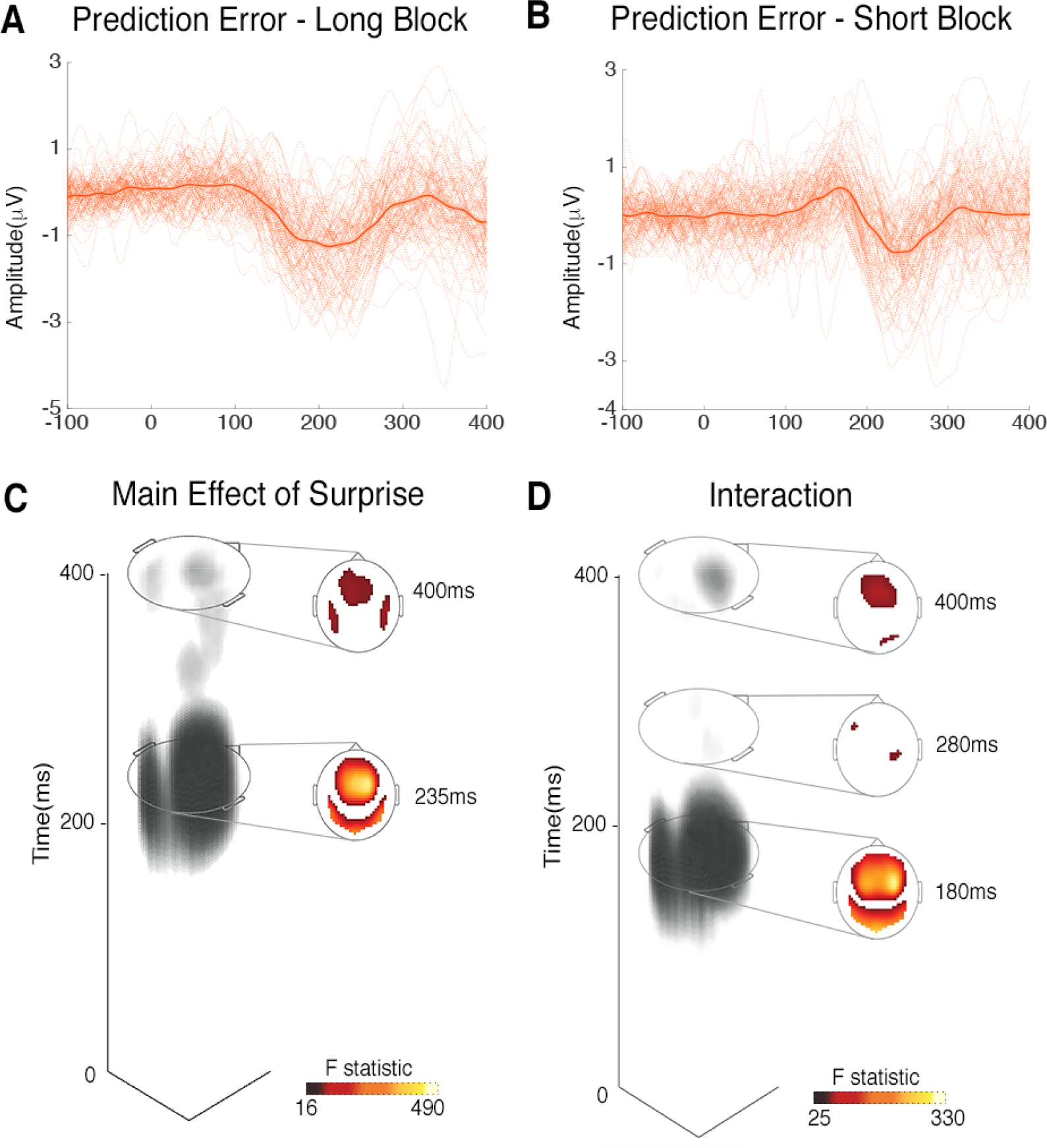
Prediction errors to classical oddball. Prediction error waveforms at the Fz channel for (A) long duration deviant block and (B) short duration deviant block for each participant (dotted lines), with grand mean across all participants (solid lines) and standard error of mean (lighter shading). Spatiotemporal statistical analysis revealed a significant (C) Main Effect of Surprise and a (D) Surprise*Deviant Type Interaction, with a greater surprise effect for long blocks. (C-D) Maps displayed at p<0.05 FWE corrected for the whole space-time volume.

A regression analysis of the prediction error response for the Long deviant block revealed a trend level negative relationship between CAPE+ scores and the long duration mismatch response [t (1,97) = 4.01, p(cluster-FWE) = 0.057] at 380ms. This is indicative of more attenuation of the mismatch response as the number of positive symptoms increases.

#### 3.1.2 Stochastic Oddball Paradigm

Prediction error waveforms (Deviants > Standards) for each variance condition (narrow and broad); (Figure 4A,4B). A spatiotemporal analysis was conducted to investigate if there was an effect of surprise and variance. An ANCOVA revealed significant main effects of *Surprise* and *Variance* as well as a significant *Surprise*Variance* interaction (Figure 4). The main effect of *Surprise* (Figure 4C) showed significant clusters over medial frontocentral channels at 150ms (*F* = 295.78, *p* < .001), left parietal channels at 145ms (*F* = 257.72, *p* < .001) and right frontocentral channels at 365ms (*F* = 92.53, *p* < .001). This is in line with previous prediction error findings in the mismatch negativity (100-250ms) and P300 (250-500ms) time-windows (Garrido et al., 2013). The *Surprise*Variance* interaction (Figure 4D) showed significant clusters over left tempo-parietal channels at 155ms (*F* = 47.85, *p* < .001), right frontocentral channels at 160ms (*F* = 41.90, *p* < .001), medial frontocentral channels at 255ms (*F* = 35.23, *p* < .001) right parieto-central channels at 155ms (*F* = 21.68, *p* = .029), and right frontal channels at 210ms (*F* = 18.15, *p* = .042). This also confirms previous findings whereby the outlier response is greater in the narrow than the broad condition (Garrido et al., 2013).

**Figure 4:**
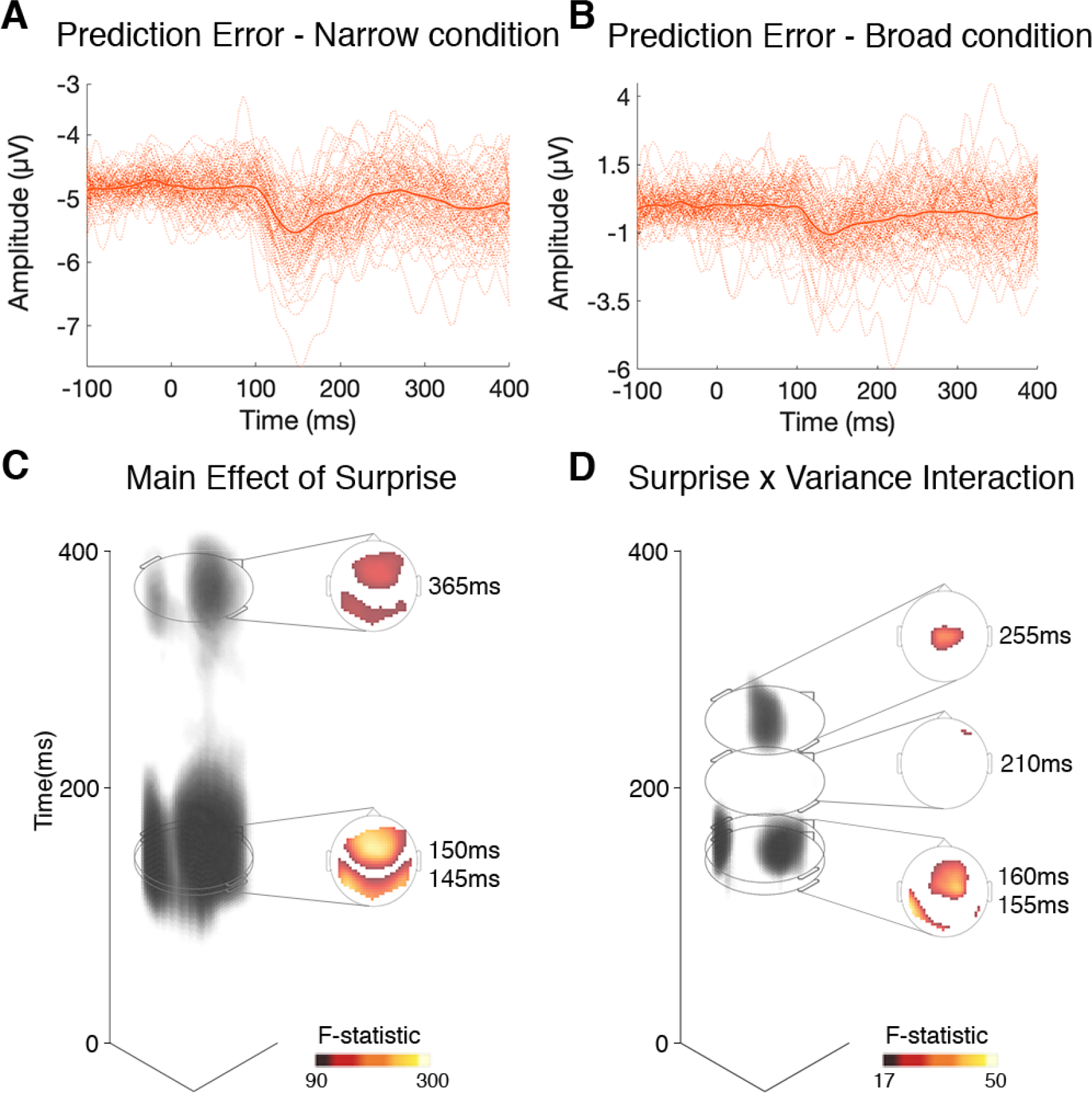
Prediction errors to stochastic oddball. Prediction error waveforms at the Fz channel for (A) narrow and (B) broad conditions for each participant (dotted lines), with grand mean across all participants (solid lines) and standard error of mean (lighter shading). Spatiotemporal statistical analysis revealed significant (C) Main Effect of Surprise and (D) Surprise*Variance Interaction.

A regression analysis with the *Surprise*Variance* interaction as outcome variable and CAPE+ scores as the predictor variable, controlling for age, medication effects and other measures of psychopathology revealed significant clusters over left tempo-parietal channels at 155ms (*t* = 5.28, *p* = .002), medial frontocentral channels at 145ms (*t* = 4.89, *p* = .007) (see Figure 4). This indicates that as the number of psychotic experiences increases, the sensitivity for learning about, and detecting violations to, statistical regularities in a stochastic environment decreases.

### 3.2 Structural and functional predictors of psychotic experiences

**Figure 5:**
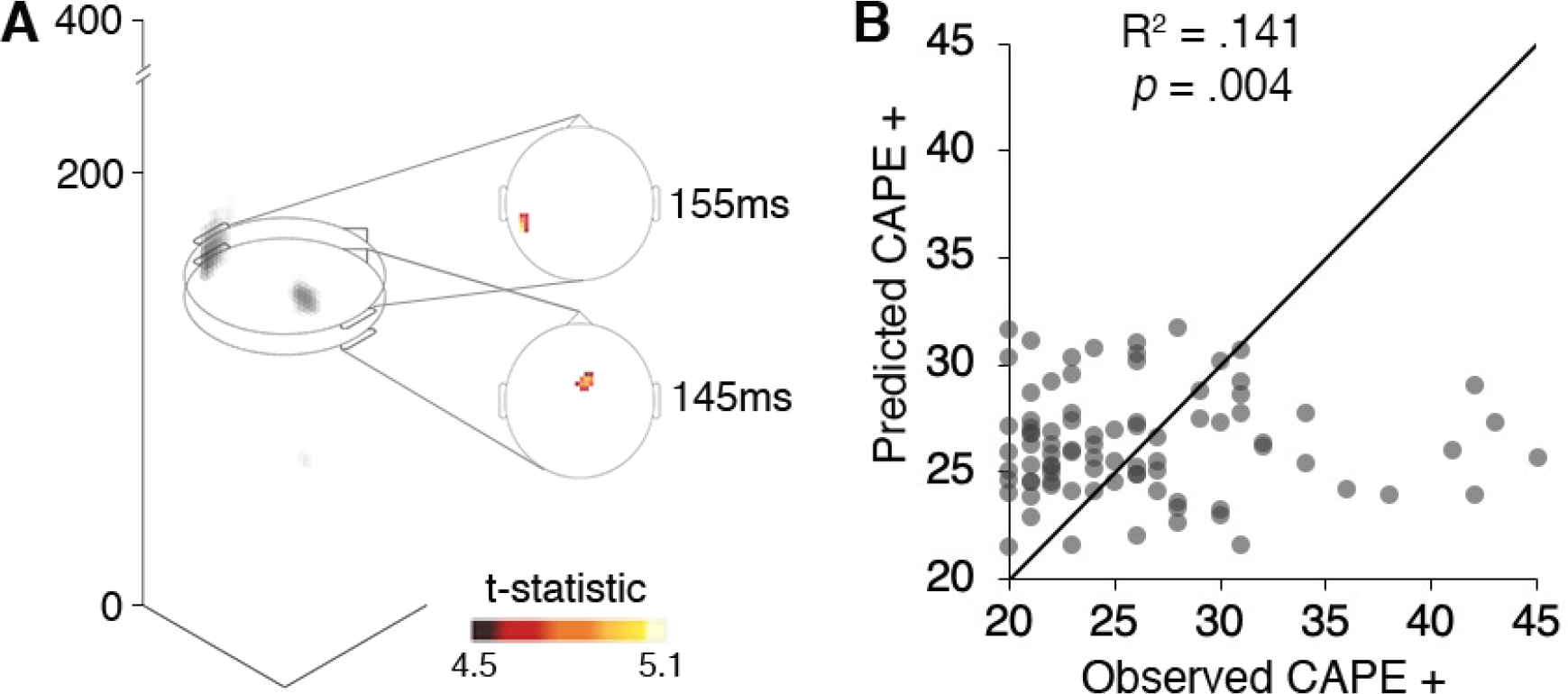
Functional and structural connectivity predict positive psychotic-like experiences. (A) A regression model found that individual prediction error responses over the two conditions were greater for smaller CAPE+ scores while controlling for age, and measures of psychopathology with significant clusters peaking at 145ms and 155ms. Maps displayed at p<0.05 FWE corrected for the whole space-time volume. (B) A regression analysis found that prediction errors (field intensity values at 145ms/155ms) and structural connectivity (FA of left arcuate fasciculus/right arcuate fasciculus/auditory interhemispheric pathway) predict CAPE+ scores. Scatterplot shows model fit between observed (actual) and predicted CAPE+ scores.

In order to investigate whether functional and structural measure together predict psychotic experiences in psychologically healthy individuals better than prediction errors alone, a multiple linear regression analysis was performed. The outcome variable was CAPE+ scores. In a first step, the prediction error amplitudes extracted from the stochastic oddball paradigm (145ms/155ms) were added to the model as predictor variables, as they were found to correlate with CAPE+. In a second step, we added FA values as predictor variables, namely the FA of the left arcuate fasciculus, the FA of the right arcuate fasciculus and the FA of the auditory interhemispheric pathway. Prediction errors alone explained 8.6% of the variance in CAPE+ scores (*F* = 5.044, adjusted R^2^ = .086, *p* = .009), but when FA of the three white matter tracts were added to the model the variance explained increased to 14.1% (*F* = 3.829, adjusted R^2^ = .141, *p* = .004). This significant increase in explained variance (*F*_change_= 2.803, R^2^ change = .084, *p* = .045), indicates that mismatch responses and structural connectivity together better predict psychotic experiences in psychologically healthy individuals (Figure 5B) than mismatch responses alone. Furthermore, FA of the auditory interhemispheric pathway (*t* = −2.694, *p* = .009) and FIVs at 145ms (*t* = −3.099, *p* = .003) were significant independent predictors of CAPE+ scores. This indicates that psychotic experiences increase as the microstructure of the auditory interhemispheric pathway and the sensitivity to statistical regularities in a stochastic environment decrease.

## 4. Discussion

The aim of the present study was to investigate functional and structural brain connectivity predictors of psychotic experiences in the healthy population. We found that reduced prediction error responses were associated with increasing positive psychotic experiences in healthy individuals. These findings indicate that as the number of psychotic experiences increases the mismatch response decreases and the ability to learn, and detect violations to, statistical regularities in a stochastic environment decreases. Critically, we found that reductions in the microstructure of the auditory interhemispheric pathway and decreasing mismatch responses predict increasing psychotic experiences in healthy individuals.

The classical oddball paradigm failed to reveal significant effects with CAPE+ scores within the MMN or P300 time windows, which is in line with Broyd et al.(2016), who did not find group differences between high and low schizotypy groups. The stochastic oddball paradigm however, found clusters that fall within the typical MMN time-window(Sams et al., 1985). This is in line with MMN attenuation consistently observed in chronic schizophrenia, first-episode psychosis, and individuals at high-risk for schizophrenia(Todd et al., 2008; Näätänen et al., 2015; Erickson et al., 2016). It should be noted that the classical and stochastic oddball paradigm differs in several ways which may explain the discrepancy of our effects in the two paradigms: 1) the deviants used (frequency vs. duration), 2) deviant probabilities (simple vs. stochastic), and 3) the cognitive load of the incidental task (i.e. one back vs. two back). Having said this, it has been shown that increasing cognitive load does not decrease prediction error amplitude (Garrido et al., 2016). Nonetheless, the finding that the stochastic, but not the classical oddball paradigm, demonstrated an association between MMN attenuation and increasing psychotic experiences suggests that complex oddball paradigms with abstract rules (like the former) may be more sensitive for revealing emerging prediction error deficits prior to any presentation of psychotic symptoms.

Here we also demonstrate that MMN amplitudes identified with the stochastic oddball paradigm in combination with DWI measures of auditory white matter microstructure are predictive of positive psychotic experiences in healthy individuals, which are more subtle than psychotic symptoms experienced by patients diagnosed with schizophrenia. It is important to note that our regression model performs well at lower CAPE+ scores but not with higher CAPE+ (35<) scores, which may be driven by the data being more skewed towards lower CAPE+ scores. However, we observed that early to mid-latency mismatch responses and the microstructure of the auditory interhemispheric pathway are independent predictors of psychotic experiences. A study investigating the association between auditory interhemispheric pathway microstructure and psychotic symptoms in schizophrenia observed decreased fractional anisotropy (indicative of loss of overall white matter microstructure) and increased radial diffusivity (indicative of reduced myelination) in patients with auditory verbal hallucinations compared to patients without auditory verbal hallucinations and healthy controls(Wigand et al., 2015). Our findings indicate that auditory interhemispheric pathway microstructure might also be associated with pre-clinical psychotic experiences and therefore represent a promising early biomarker of psychosis.

Contrary to findings in schizophrenia whereby reduced microstructure of the arcuate fasciculus has recurrently been linked to psychotic symptoms such as auditory verbal hallucinations(Geoffroy et al., 2014; McCarthy-Jones et al., 2015), we did not observe the arcuate fasciculi to be independent predictors of psychotic experiences in healthy individuals. White matter microstructure has been reported to deteriorate gradually with illness duration from early to chronic stages of schizophrenia(Friedman et al., 2008; Di Biase et al., 2017) and previous studies failed to observe microstructural changes of the arcuate fasciculus in first-episode schizophrenia(Peters et al., 2008) or individuals at high-risk for schizophrenia(Munoz Maniega et al., 2008). It is therefore possible that microstructural changes associated with psychotic experiences are initially localized to auditory pathways in the corpus callosum and then advance to association tracts like the arcuate fasciculus with symptom presentation and prolonged illness duration.

The lack of clinical schizophrenia groups limits the extent to which the findings of the present study might be informative for the identification of psychosis biomarkers. Future studies including such cohorts will be key in determining whether the structural and functional predictors of *psychotic experiences* identified in the present study are valuable *psychosis symptom* predictors in individuals at high-risk for psychosis, first-episode schizophrenia patients and chronic schizophrenia patients. Further, longitudinal studies will be critical in elucidating whether these structural and functional individual brain alterations can provide prognostic clues about illness progression.

In conclusion, we found that individual decreases in the microstructure of the auditory interhemispheric pathway and reduced sensitivity to auditory regularities in a stochastic environment predict psychotic experiences in healthy individuals. To the extent that these structural and functional brain connectivity changes have previously been reported in patients with schizophrenia, the findings of this study suggest that both MMN and the microstructure of the auditory interhemispheric pathway may have a translating potential as targets for early screening of psychosis.

## Acknowledgements

We thank John McGrath for helpful discussions. David Lloyd, Elise Rowe for support in EEG data acquisition. Aiman Al-Najjar and Nicole Atcheson for conducting MRI data collection, and all participants for their time. M.I.G. was supported by a UQ Fellowship (2016000071), a UQ Foundation Research Excellence Award (2016001844), and the ARC Centre of Excellence for Integrative Brain Function (ARC CE140100007). R.R. was supported through a UQ International Research Scholarship.

